# Diversity and Functional Roles of Carabid Beetles across Salinity Gradients in Marshlands

**DOI:** 10.64898/2025.12.04.692354

**Authors:** S. Remmers, K.H. Dausmann

**Affiliations:** Functinal Ecology, Institute of Cell and Systems Biology of Animals, Universität Hamburg, Germany

**Keywords:** marsh ecosystems, carabid beetles, estuary, salinity, biodiversity

## Abstract

Marsh ecosystems represent ecotones of high ecological significance where terrestrial and aquatic systems converge, governed by strong gradients in salinity and flooding. These environments sustain specialised communities and perform vital ecosystem functions such as carbon sequestration or nutrient cycling. This study examined carabid beetle (Coleoptera: Carabidae) assemblages across freshwater and saltwater marshes along the Elbe estuary in northern Germany to assess how environmental conditions shape patterns of abundance, diversity, and functional traits. Over 17,000 individuals representing 84 species were collected between March and October 2024. Despite similar species richness between marsh types, community composition differed markedly (Bray–Curtis dissimilarity: 0.92), reflecting environmental filtering along the salinity gradient. Freshwater marshes exhibited greater overall functional richness, while saltwater marshes supported more functionally even and divergent trait distributions, indicating stronger niche specialization under saline stress. Seasonal activity was bimodal in both habitats but peaked earlier in freshwater sites. A generalized linear mixed model revealed significant positive relationships between carabid activity abundance and functional richness of both dispersal- and preference-based traits. These findings demonstrate that salinity and flooding regimes drive distinct taxonomic and functional assemblages, with trait diversity underpinning ecosystem resilience and functioning. The study establishes an ecological baseline for future monitoring of marshland biodiversity and highlights the necessity of conserving functionally diverse communities to maintain the ecological integrity of tidal and floodplain marshes under changing environmental conditions.

## Introduction

Patterns of biodiversity and species abundance provide essential insights into ecosystem functioning and trophic interactions. These patterns can be particularly pronounced in marsh ecosystems, as ecotones where terrestrial and aquatic systems converge (Lambeets et al., 2008). Tidal marshes and estuarine systems are important for the preservation of ecological processes and biodiversity and offer critical ecosystem services such as flood protection, carbon sequestration, and nutrient cycling. These ecosystem services are directly linked to the health of key species and their interactions with the ecosystem, supporting complex food webs (Bhuiyan et al., 2025; Colombano et al., 2021). Their waterlogged soils are rich in organic carbon due to slow decomposition, making them important carbon sinks that contribute to climate mitigation (Huyzentruyt et al., 2025; Ouyang & Lee, 2014). Despite high productivity, they support a limited number of specialized terrestrial and marine species adapted to fluctuating salinity and flooding (Greenberg et al., 2006). Estuaries, located at the transition between terrestrial and marine environments and where freshwater from rivers meets saltwater from the sea, create distinct hydrological zones. Near the mouth of the river, salt marshes with high salinity and frequent tidal flooding are found, whereas further upstream freshwater marshes characterized by more riverine input and stable conditions occur. These variations influence soil properties, nutrient availability, and species community structure, shaping the ecology and biodiversity of the entire system (Åhlén, 2024).

Within these dynamic habitats, carabid beetles (Coleoptera: Carabidae) are excellent bioindicators for assessing ecosystem condition as well as investigating ecosystem interactions and cross-boundary connectivity (Koivula, 2011; Riley Peterson et al., 2021). Carabid beetles represent one of the largest and ecologically most important families within the order Coleoptera, known for their remarkable morphological and ecological plasticity (Lövei & Sunderland, 1996; Nolte, 2018). They play a key role in ecosystems by acting as invertebrate predators, feeding on potential pests, and consuming weed seeds (Müller et al., 2022; Ali & Willenborg, 2021). Carabid beetles vary greatly in size, colouration and anatomical adaptations such as specialized digging tibiae and wing dimorphism, traits that reflect their ground-dwelling lifestyles and contribute to their evolutionary success (Aukema, 1995; Thiele, 1977). Additionally, they serve as prey for other animals, and their varied diet helps regulate insect populations, which can indirectly support plant health (Bonacci, 2025; De Heij & Willenborg, 2020). The diversity in their traits forms the basis of their adaptability across various habitats and ecological niches. In Germany alone, approximately 580 species are known, many of which are habitat specialists with diverse ecological requirements (Schmidt et al., 2016; Trautner et al., 2014). There is considerable interspecific variation in habitat preferences, ranging from stenotopic species adapted to narrow environmental conditions to generalists (eurytopic species) capable of tolerating abroad range of environments. The presence or absence of particular species, especially specialists, provides valuable insight into habitat characteristics and changes in local environmental conditions (Nagel, 1997). Because of their diversity, sensitivity and ecological functions, carabids are widely recognized as bioindicators for specific habitat conditions such as soil structure, nutrient and moisture content and ground-level vegetation (Dornieden et al., 2005). Their high mobility and dispersal capacity enable them to respond more quickly to environmental changes than, for example, vegetation, making them particularly well-suited for detecting disturbances or shifts in habitat quality and rapid anthropogenic changes (Malheiros et al., 2025; Rainio & Niemelä, 2003; Trautner & Aßmann, 1998).

Despite their ecological importance and the complex interplay of marshland hydrology and species adaptation, most research to date has concentrated on agricultural system, driven largely by economic interests and the relevance of pest control and landscape management. However, understanding the ecological dynamics of natural habitats like tidal marshes is crucial, as these areas contribute substantially to biodiversity and ecosystem functioning, yet remain comparatively understudied. Investigations in natural marshes not only enhance our understanding of fundamental ecosystem processes but also provide reference points for evaluating the impacts of anthropogenic environmental changes, assessing the effectiveness of conservation and for understanding the resilience and vulnerability of different ecosystems under environmental change (Alemu I et al., 2024; Fukami & Wardle, 2005). Establishing ecological baselines in relatively undisturbed marshlands allows for more accurate assessments of shifts in habitat conditions in altered or managed landscapes, including agricultural fields and restored wetlands. To gain a more comprehensive understanding of ecosystem functioning, it is necessary to expand research to include the functional ecology of characteristic species such as carabid beetles, which play key roles in marshland food webs and trophic dynamics.

The Elbe River forms an extensive tidal estuary of saltwater, brackish, and freshwater marshes, with gradients in salinity and tidal influence. Stretching from the open North Sea at Cuxhaven far inland to the Geesthacht weir, the estuary is bordered by dikes along most of its length. These dikes create low-lying floodplain areas and provide a diverse range of habitats with significant ecological and conservation value (Elbe Estuary Working Group, 2012).

This study investigated how contrasting environmental conditions in saltwater and freshwater marshes along the Elbe estuary shape the abundance, biodiversity and functional diversity with its ecological roles of carabid beetles. By analysing carabid communities from paired saltwater and freshwater marshes, we aimed to gain new insights into their species richness and habitat affinities, thereby improving our understanding of their functional roles in these dynamic ecosystems. Furthermore, we assessed seasonal variations in beetle activity and examined species composition along hydrological gradients from low to high marsh zones, considering different flooding regimes. By documenting these patterns, our work establishes an ecological baseline for future studies of food web structure, nutrient transfer, and the functional role of carabid beetles in marshland carbon and energy cycles.

## Methods

This study was conducted in the Elbe estuary in Northern Germany (Fig. 1). We studied two sites along the estuary’s salinity gradient: a freshwater marsh (53.6619°N / 9.5513°E) and a saltwater marsh (53.9325°N / 8.9121°E). The freshwater marsh is part of the Haseldorfer Binnenelbe nature conservation area, while the saltwater marsh is located within the Schleswig-Holstein Wadden Sea National Park. The different marsh types are characterized by different levels of salinity. In 2021, soil salinity in the high marshes at a depth of 0-4 cm was measured at 0.51 ‰ in the freshwater marsh and 6.82 ‰ in the saltwater marsh (Lexmond, 2025).

**Figure 1:**
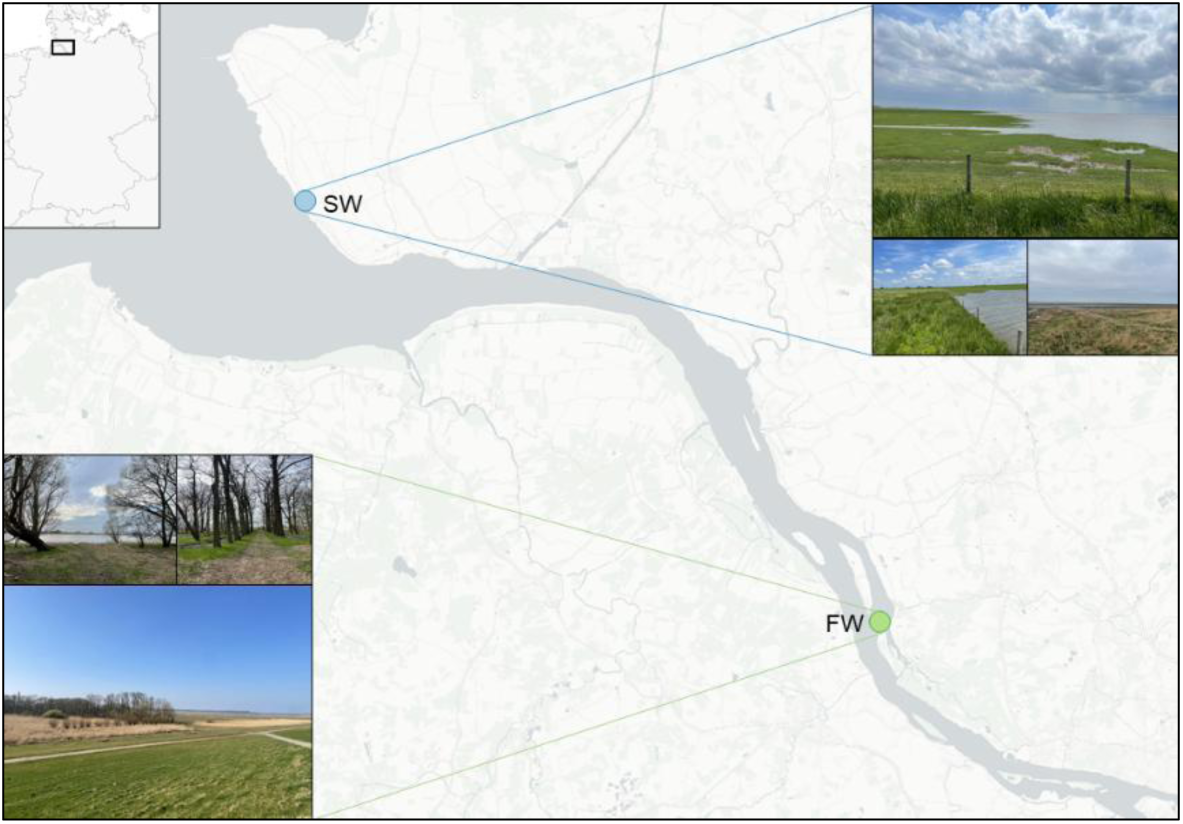
Map showing the locations of the study sites in the freshwater marsh (FW; green) and the saltwater marsh (SW; blue). Photographs depict the characteristic habitats of each marsh type.

The freshwater marsh is characterized by floodplain meadows, with patches dominated by slender or riparian sedges and reeds (e.g., *Agrostis stolonifera, Phragmites australis, Juncus effuses, Carex hirta*; LSH Biotopbogen). Because one of the transects in the freshwater marsh was located along a tree-lined avenue/path within an alluvial forest patch, a characteristic element of this river floodplain nature conservation area, other types of vegetation were also dominant in this marsh, such as basket willows (*Salix viminalis*), alders (*Alnus glutinosa*), cleavers (*Galium aparine*) and stinging nettles (*Urtica dioica*). The saltwater marsh is an extensive coastal wetland with tidal creeks that transitions directly into adjacent mudflats. The ungrazed area is dominated by halophilic vegetation, such as rushes, fescue grasses, glassworts and cordgrasses (e.g. *Elymus spp., Festuca rubra, Puccinellia maritima, Spartina anglica, Salicornia spp.*). Both marsh types are grazed by sheep; however, all transects were located in ungrazed areas of the marshes.

Within each marsh, two transects were established by installing pitfall traps orthogonally to the Elbe River, extending from the boundaries of the low marsh (mean high water line) to the high marsh to account for variation along the flooding regime and distance from the riverbank. (Fig. 1). Low marshes are typically flooded twice a month during spring tides, whereas high marshes are flooded only a few times per year, usually during storm tides.

In total, 42 pitfall traps were installed along four transects and left open from March to October 2024. Transects F1 (11 traps) and F2 (9 traps) were located in the freshwater marsh, while transects S3 (10 traps) and S4 (12 traps) were located in the saltwater marsh of the Elbe estuary (Fig. 1). Pitfall traps were installed at intervals of 25 m, resulting in transect lengths of 250-300 m from the riverbank inland, except for transect S3, where the distance between traps was reduced to 10 m due to the limited marsh area in front of the dike (Fig. 1). This trapping technique (Barber, 1931) is considered the standard method for recording ground living arthropods, like carabid beetles (Brown & Matthews, 2016; Trautner, 1992) (Fig. 2). Pitfall traps were plastic cups that were buried with the upper edge of the trap being level with the ground surface (opening diameter: 9cm; height: 12 cm). The traps were filled with Renner solution (Renner, 1982; 40% ethanol, 35% water, 20% glycerol, 5% acetic acid and a drop of dishwashing detergent to reduce the surface tension) as a preservative, and covered with wire mesh (mesh size: 1-1.5 cm) to prevent vertebrates from falling in and to minimize the accumulation of leaf litter. Traps were checked regularly and emptied every 14 days, with more frequent checks during periods of high temperatures or after heavy rainfall to avoid evaporation or dilution of trapping solution. Traps were labelled in the field and stored at 7 °C until sample processing.

**Figure 2:**
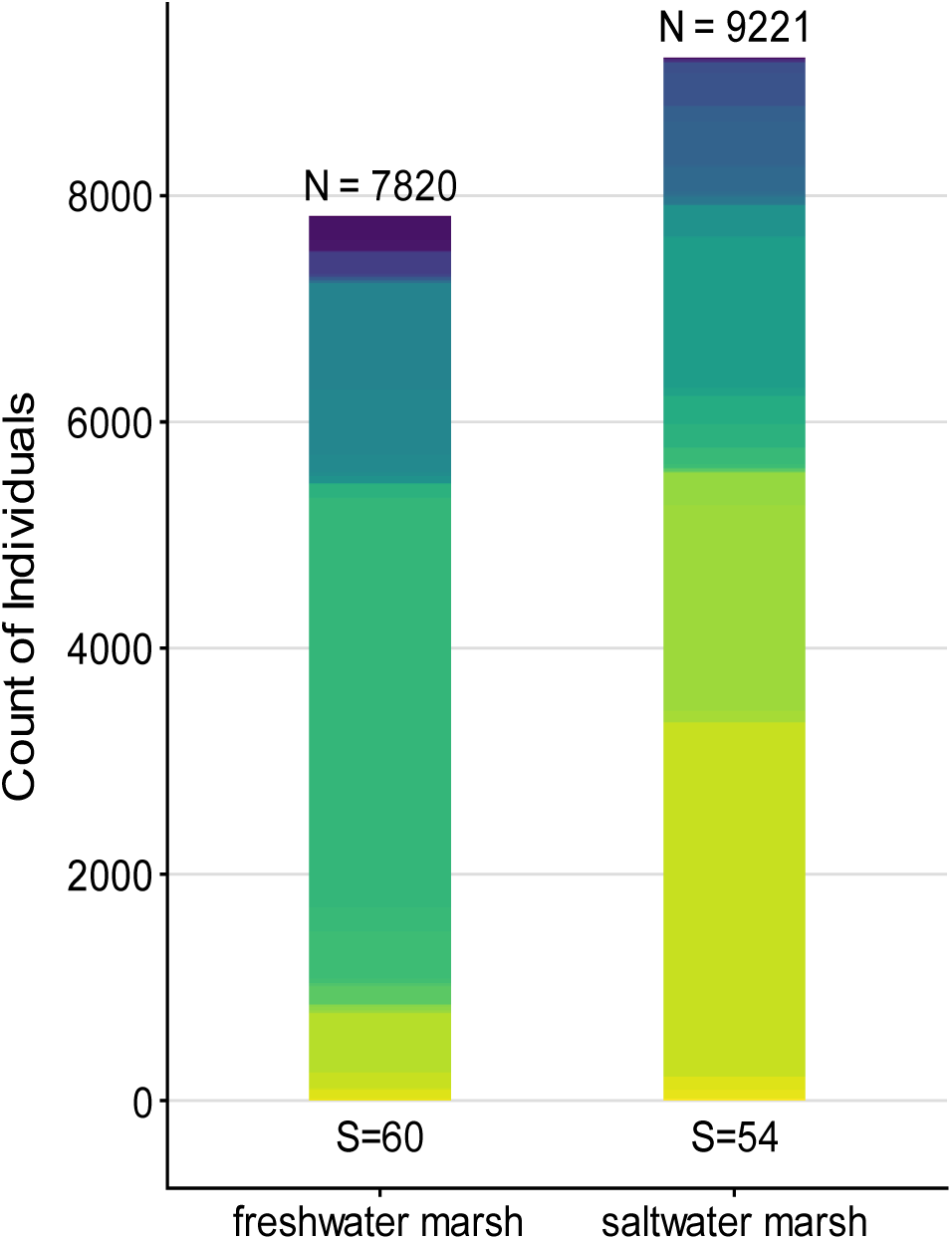
Abundance of carabid beetles in the freshwater and saltwater marshes. “N” indicates the total number of individuals, while “S” shows the number of species in each marsh type. Different colors represent the distinct carabid beetle species.

### Taxonomy and Sample Processing

A total of 719 traps were processed, with 207 traps from transect F1, 133 traps from F2, 164 traps from S3 and 215 traps from S4, excluding those that could not be collected due to heavy flooding or destruction. In the laboratory, the contents of the pitfall traps were sorted into carabid beetles, spiders, and other animal groups (bycatch). Carabid beetles were sorted using a stereo microscope (Leica EZ4 16/15, maximum optical magnification 56x), identified to species level and their sex was determined (Freude et al., 1976; Lompe, 2025; Müller-Motzfeld, 2012). Taxonomic identifications were confirmed during carabid beetle conferences, with some rare species additionally validated by experts. Carabid beetle specimens from the same trap were preserved together in vials filled with 96% ethanol.

Species habitat preferences were classified according to the Catalogue of Habitat Preferences (GAC e. V., 2009). Additionally, the habitat breadth or each species was calculated. The definition and grouping of habitat types is further explained in the Supplementary Materials (*Habitat Types and Preferences*). To quantify different facets of functional diversity in ecological communities beyond taxonomic diversity and abundance, we assigned ecological traits to species i.e., body size, wing morphology, moisture preference, halophily status, as well as strata and trophic level (Klaiber et al., 2017). To account for differences in sampling effort, activity abundance was calculated for each pitfall trap and species. This was done by first summing the number of individuals collected in each trap, dividing it by the number of days the trap was active and then scaling this value to a 14-day trapping period. This yielded an activity abundance measure standardized to two weeks. For each species within each marsh, the mean activity abundance and standard deviation were then calculated across all traps.

### Statistical Analyses

The statistical tests applied to each analysis are specified directly within the Results section alongside the relevant findings.

### Estimator

In ecological surveys, usually not all species actually present are detected. To estimate the true number of species present, we applied the abundance-based species richness estimators “Chao1” and “ACE” (Abundance Coverage Estimator) to the combined species counts from individual traps per marsh (Chiu et al., 2014). These estimators helped assess sampling completeness by estimating how many species were detected relative to those likely present. Further methodological details are provided in the Supplementary Materials (*Species Richness Estimator)*.

### Diversity Indices

Shannon diversity was calculated for each sample (traps and temporal replicas) to measure species diversity and incorporates both the number of species present and evenness. A non-metric multidimensional scaling (NMDS) analysis based on Bray-Curtis dissimilarity was performed using species abundance data to quantify differences in carabid beetle assemblages between and within the two marsh types (Leyer & Wesche, 2008).

### Functional diversity

We generated two complementary sets of ecological traits for further analysis. The first set consisted of continuous and categorical traits relevant to dispersal and habitat specialization (body size, habitat breadth, wing morphology), hereafter referred to as dispersal traits. The second set included only categorical traits related to ecological preferences (halophily status, moisture preference, strata and trophic level), hereafter referred to as preference traits. We calculated functional diversity indices from these trait sets, Functional Richness (FRic), Functional Evenness (FEve), Functional Divergence (FDiv) and Functional Dispersion (FDis). FRic reflects the potential functional trait capacity of a community, regardless of species abundances. FEve quantifies how evenly abundances fill the occupied trait space. FDiv quantifies niche differentiation and resource use based on the distribution in trait space and was only calculated for the dispersal traits. FDis, reflecting the functional dispersion or diversity of the community, was calculated only for the preference traits, as FDiv cannot be applied to categorical data. Both FEve and FDiv values always range between 0 and 1, while FRic and FDis values can exceed 1, depending on the scale of the trait space. Calculations were based on Villéger et al. (2008), which compute multidimensional indices using a distance-based principal coordinates analysis (PCoA; (Laliberté & Legendre, 2010).

### Model

A generalized linear mixed model (GLMM) with a negative binomial distribution (count data and overdispersion) was used to analyze the factors influencing activity abundance, incorporating both fixed and random effects. The full model included the predictors marsh, month, distance from riverbank, mean body size and functional diversity indices for both trait sets. Model simplification was performed by backward selection, with non-significant (p > 0.05) predictors dropped if both AIC and likelihood ration tests suggested no substantial loss of model fit, resulting in a final model with month, marsh type, mean body size, functional richness for both trait sets and functional evenness for dispersal traits as fixed effects and species and trap as random intercepts. Model diagnostics included checks for overdispersion and inspection of residual plots to assess model fit. All analyses were conducted in R version 4.5.1 using the glmmTMB and DHARMa packages (R Core Team, 2025; Posit team, 2025; Brooks et al., 2017; Hartig, 2016).

## Results

### Species Richness and Estimators

In total, we sampled 17,039 carabid beetles belonging to 84 species (Gamma diversity) in the Elbe estuary. Of these, 7,820 individuals from 60 species were found in the Freshwater marsh and 9,221 individuals from 54 species were found in the saltwater marsh, showing a similar species richness for both marsh types (Fig. 2). We identified 26 species represented by fewer than five individuals, and 12 out of the total 84 species are classified as threatened, according to the Red List of Carabid Beetles in Germany (Schmid et al., 2016).

We generated a sample-based rarefaction curve, representing a statistically expected species accumulation curve. This approach allowed us to visualize species richness in relation to sampling effort (Achtziger, Nigmann & Zwölfer, 1992) (Supplementary Materials, Fig. I). Additionally, estimates of species richness based on ecological estimators were calculated. The results from both Chao1 and ACE estimators suggest that our sampling effort captured the majority of the estimated species richness in the saltwater marsh (96% and 93%, respectively), while in the freshwater marsh, sampling covered approximately 79 % of the estimated species richness (Tab. 1).

**Table 1:** Species richness estimates for carabid beetles in the freshwater and saltwater marshes. Observed species number (S_obs_), Chao1 and ACE richness estimators (± standard error), and the percentage of observed species relative to each estimator.

|  | freshwater marsh | saltwater marsh |
| --- | --- | --- |
| $S_{obs}$ | 60 | 54 |
| $Chao1 \pm se$ | $75.17 \pm 10.85$ | $55.87 \pm 2.26$ |
| $\% S_{obs} / Chao1$ | 79.82 | 96.64 |
| $ACE \pm se$ | $75.47 \pm 4.26$ | $57.87 \pm 3.71$ |
| $\% S_{obs} / ACE$ | 79.50 | 93.30 |

### Seasonal Activity Shifts

Both marsh types showed clear seasonal variation in carabid activity abundance. However, the timing of peak activity differed. The freshwater marsh reached its highest values earlier, in spring and early autumn, whereas the saltwater marsh peaked later, in summer (Fig. 3).

**Figure 3:**
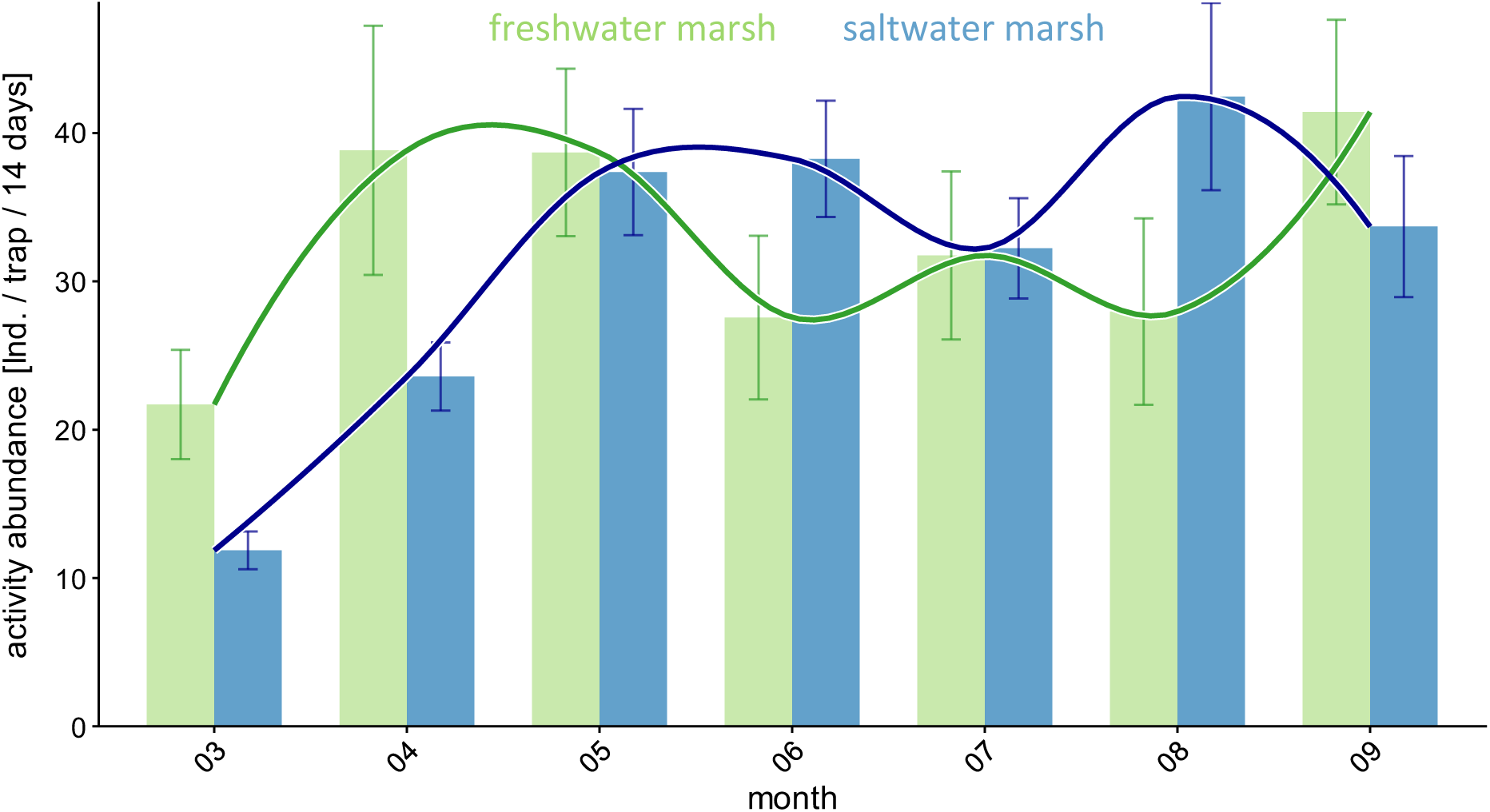
Seasonal patterns of carabid beetle activity abundance in the Elbe estuary throughout the annual activity period. Bars represent the mean activity abundance for each marsh type (freshwater marsh: green; saltwater marsh: blue) and month, with error bars indicating standard error. The overlaid lines represent LOESS curves, which use locally weighted regression to reveal non-linear trends across the months for each marsh type.

In the freshwater marsh carabid activity abundance was lowest in March at the onset of the beetles’ activity period (21.7 ± 3.7 individuals/trap/14 days) and reached a pronounced peak in April and May (38.8 ± 8.4 and 38.7 ± 5.6 individuals/trap/14 days, respectively). Activity declined during early summer but increased again in September (41.4 ± 6.2 individuals/trap/14 days), resulting in two notable peaks within the year. In the saltwater marsh, the overall trend was also characterized by two activity peaks, but with a more gradual seasonal development. Activity abundance was initially low in March (11.9 ± 1.3 individuals/trap/14 days), with the first increase observed in May and June (37.4 ± 4.3 and 38.3 ± 3.9 individuals/trap/14 days, respectively) and the highest values recorded in August (42.5 ± 6.3 individuals/trap/14 days), after a small decrease in July.

### Diversity Indices and Abundances

Shannon diversity displayed distinct patterns along the distance-from-riverbank gradient in freshwater and saltwater marshes (Fig. 4). Across both marsh types, Shannon diversity tended to be higher at intermediate distances from the riverbank. In the freshwater marsh, mean Shannon diversity was relatively low at the closest distances (0–20 m; ∼0.4–0.7) but increased to a peak at intermediate distances (50–70 m; ∼1–1.2) and remaining above 0.8 for most further points. The saltwater marsh showed a similar pattern, with lower diversity near the water (35–55 m; ∼0.5–0.8), followed by a pronounced peak at intermediate distances (115–175 m; ∼1.2–1.7), and a gradual decline at greater distances (>200 m).

**Figure 4:**
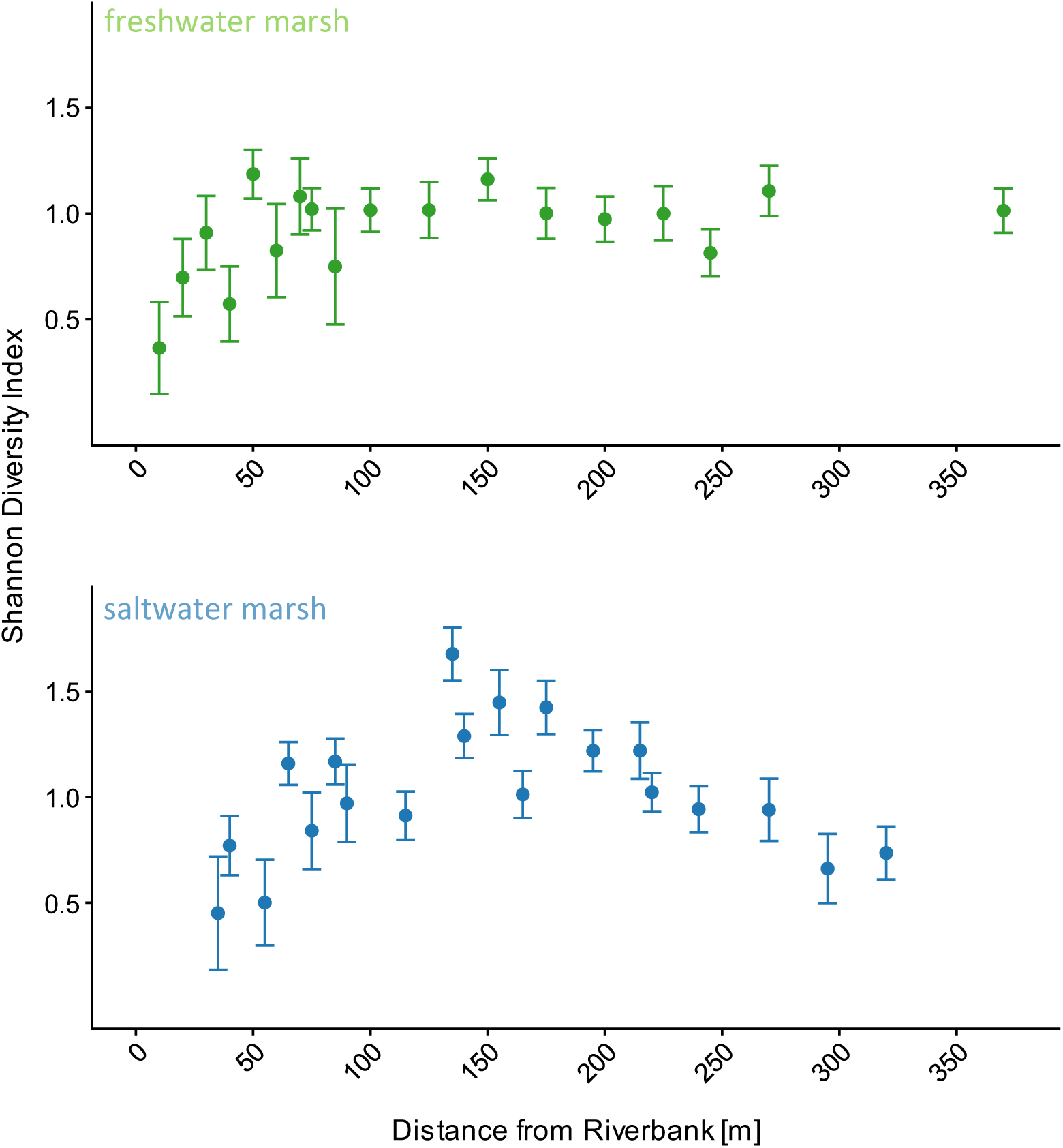
Variation in mean Shannon Diversity (± se) of carabid beetle assemblages along the distance-from-riverbank gradient in the freshwater (green) and saltwater marsh (blue). Points represent the mean Shannon diversity for traps (rounded to nearest 5 m), with error bars indicating standard error. Data from transects were pooled within each marsh type to visualize general patterns.

The beta diversity analyses revealed strong differences in community composition both within and between marshlands. In the NMDS ordination, samples from freshwater and saltwater marshes formed distinct clusters, reflecting divergent species compositions (Fig. 5). The calculated beta diversity dissimilarity values among freshwater transects reached 0.78, and saltwater transects exhibited a value of 0.79, indicating a substantial species turnover within the same habitat type in both marshes. Comparisons between marsh types showed an even higher dissimilarity of 0.92. Community composition was significant different between marsh types (PERMANOVA; R² = 0.13, F = 101.8, p < 0.001) and among sites (PERMANOVA; R² = 0.22, F = 62.13, p < 0.001). Additionally, a test for multivariate homogeneity of group dispersions (F = 18.68, p < 0.001) indicated that considerable within-site variation contributes to the observed differences among sites. Together, these results demonstrate that freshwater and saltwater marshes support distinctly different carabid beetle communities, with a high degree of species turnover among transects.

**Figure 5:**
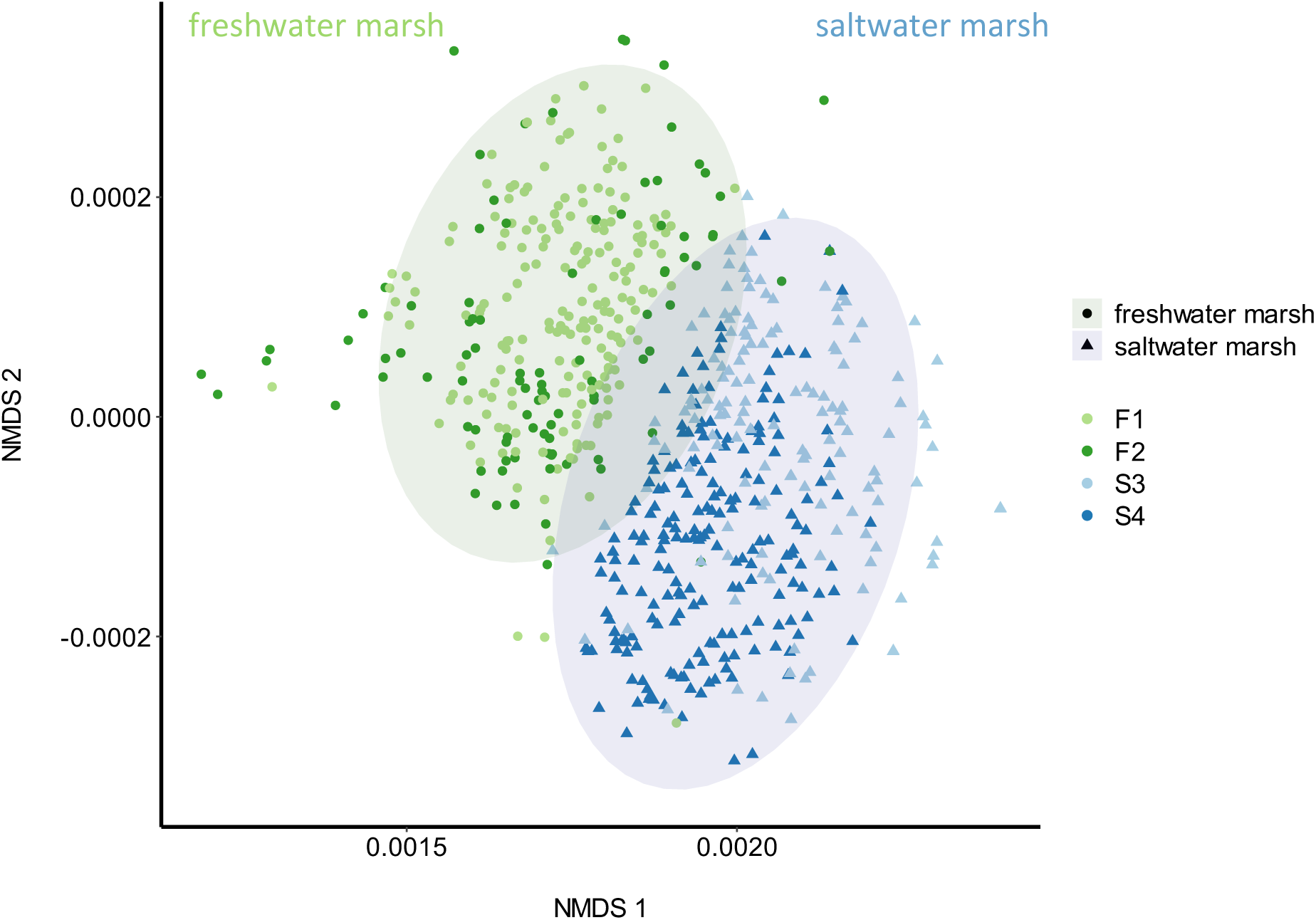
Non-metric multidimensional scaling (NMDS) plot based on Bray-Curtis dissimilarity (stress = 0.038). Each point represents a trap sample, positioned according to relative similarity in carabid beetle community composition. The axes are arbitrary and do not represent specific gradients.

Distribution of the 20 most common species, each occurring with more than 50 individuals per site, revealed clear differences in zonation patterns along the transects between freshwater and saltwater marshes (Fig. 6). In the F1 transect in the freshwater marsh, the highest relative abundances for most species were concentrated close to the riverbank (∼25-75 m; Fig. 6 F1), with abundances generally decreasing as distance from the Elbe River increased. The F2 transect extended inland for only about 100 m from the riverbank, due to the limited marsh area in front of the dike. Most traps were located within the low marsh zone and an extensive high marsh was absent, so no clear statements about zonation patterns can be made for this transect. Nevertheless, relative abundance was noticeably highest near the riverbank (∼25 m; Fig. 6 F2) and declined visibly over a short distance. In contrast, in the saltwater marsh the carabid assemblages were less concentrated at the riverbank and tend to be more evenly distributed or showed peaks at intermediate distances. Many of the most abundant species had their highest relative abundances further inland (∼125-175 m; Fig. 6 S3 and S4) and fewer species dominating close to the riverbank.

**Figure 6:**
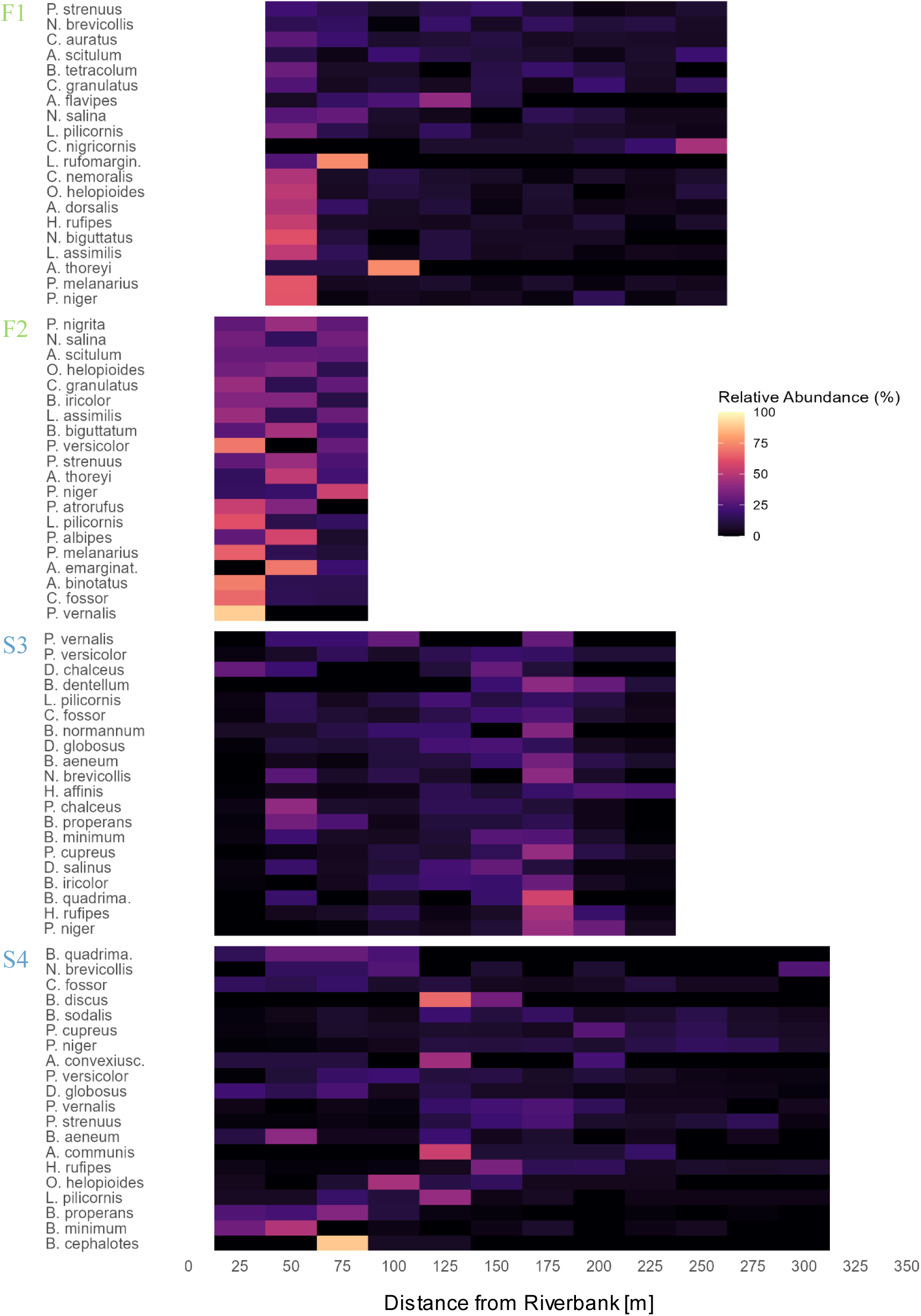
Heatmaps showing the relative abundance (%) of the 20 most common carabid beetle species along the distance-from-riverbank gradient at each site. Species are ordered from the most to least evenly distributed based on species evenness calculations. The x-axis shows distance from the water’s edge (in meters). Each bin represents a trap, rounded to the nearest 25 meters. Relative abundance values are log-transformed to better visualize both dominant and less common species within each site.

### Habitat Preferences and Functional Trait Diversity

The recorded species demonstrated a strong association with wetland environments, but also revealed considerable overlap and habitat generalism among several species. Of the 61 carabid beetle species recorded in the freshwater marsh, the majority (n = 58) had habitat preferences for coastal shores, banks, and wetland habitats. Seventeen species were classified as eurytopic, reflecting their broad occurrence across multiple habitat types. Several species exhibited overlapping habitat preferences, particularly between coastal shores, banks, and wetland habitats and either forests and woodlands (n = 15) or open, dry, or cultivated biotopes (n = 19) (see Venn diagram, Fig. 7 A). Similarly, in the saltwater marsh, most of the 58 carabid beetle species were also associated with coastal shores, banks, and wetland habitats (n = 56). Fewer species preferred forests and woodlands (n = 7), while the number preferring cultivated or dry, open biotopes and the number of eurytopic species were comparable to those in the freshwater marsh (Fig. 7 B).

**Figure 7:**
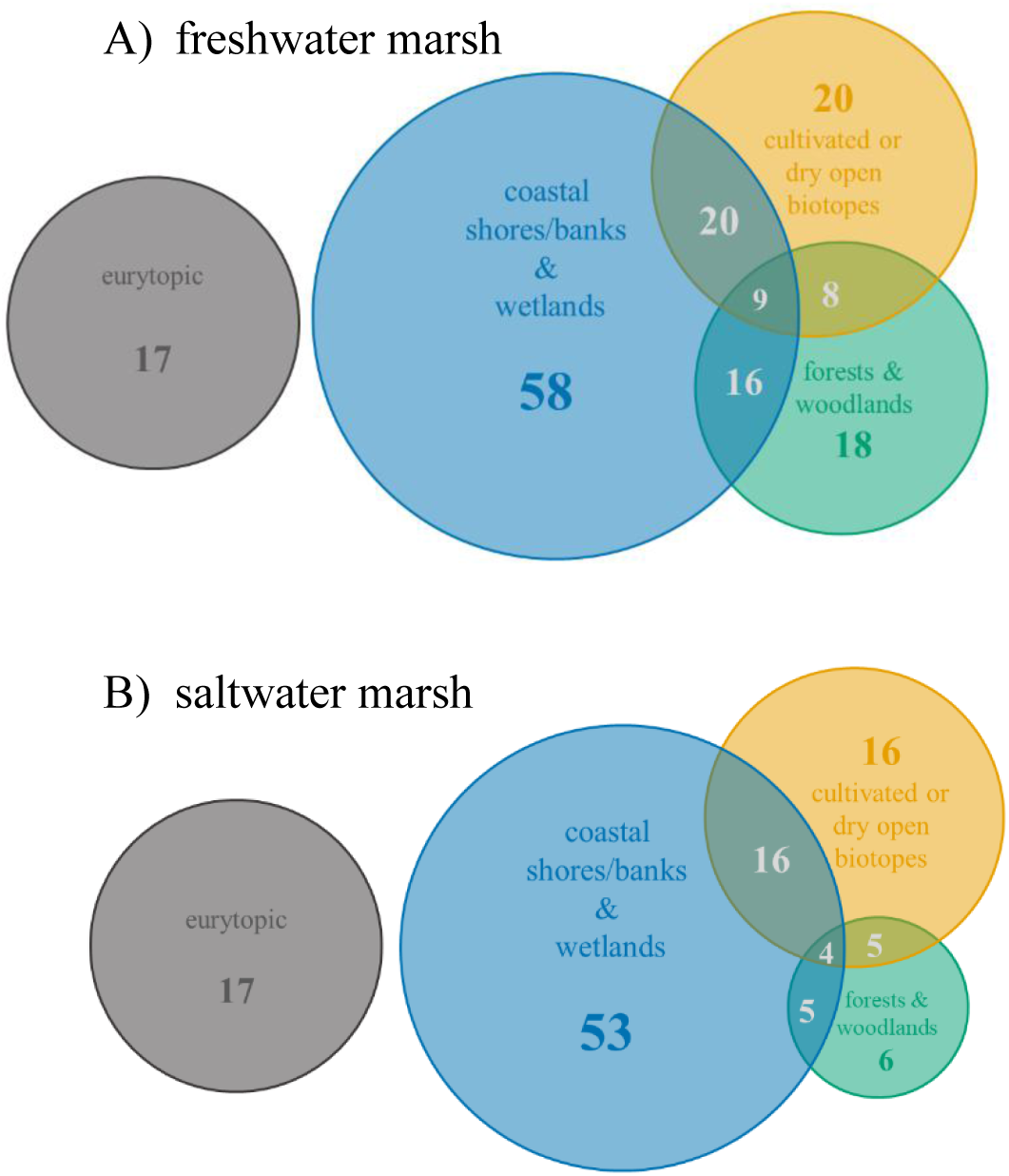
Venn diagram with four colored circles representing the habitat preferences of the carabid beetle species recorded in the freshwater marsh (A) and the saltwater marsh (B). Numbers in the non-overlapping sections indicate the total number of species that prefer each habitat type (among others; no ranking). Numbers in the overlapping sections show the number of species preferring two or three habitat types. Species occurring in more than three habitat types were classified as eurytopic

The results of the functional diversity indices for each transect are summarised in Table 2. For the dispersal traits, the freshwater transects showed higher functional richness (FRic_1_), especially at F1, indicating a broader range of dispersal traits among carabids compared to saltwater sites. However, saltwater transects had higher divergence (FDiv_1_) and similar functional evenness (FEve1), suggesting a narrower trait range that is evenly distributed, possibly reflecting greater specialization or niche differentiation. For preference traits, saltwater transects exhibited the highest functional richness (FRic_2_), evenness (FEve_2_) and dispersion (FDis_2_), indicating a more diverse and evenly distributed set of trait combinations Overall, trait combinations in saltwater marshes were less dominated by a few types, and the higher FDis_2_ implies greater functional differentiation, highlighting a broader spectrum of ecological strategies among carabid beetles.

**Table 2:** Functional diversity indices for each transect, calculated for two separate trait sets. Trait set 1 includes mixed continuous and categorical dispersal traits, while trait set 2 includes only categorical preference traits.

|  | **F1** | **F2** | **S3** | **S4** |  |
| --- | --- | --- | --- | --- | --- |
| $FRic_1$ | 0.102 | 0.072 | 0.095 | 0.093 | } |
| $FEve_1$ | 0.581 | 0.574 | 0.579 | 0.519 | |
| $FDiv_1$ | 0.761 | 0.800 | 0.841 | 0.906 | |
| $FRic_2$ | 6.25 | 7.78 | 9.23 | 9.76 | } |
| $FEve_2$ | 0.200 | 0.285 | 0.374 | 0.341 | |
| $FDis_2$ | 0.109 | 0.240 | 0.330 | 0.254 | |

Dispersal traits:
- body size - habitat breadth - wing morphology
Preference traits:
- halophily status - moisture preference - strata - trophic level

There were clear differences in trait distributions between the freshwater and saltwater transects (Fig. 8). The only statistically significant differences in the number of carabid species inhabiting the two marsh types were observed for halophily status (Fisher’s exact test, p < 0.01) and vertical strata (Fisher’s exact test, p = 0.01). A Wilcoxon rank-sum test further revealed a significant difference in body size, with carabid beetles from the freshwater marsh exhibiting larger body sizes than those from the saltwater marsh (W = 5009, p = 0.031; Supplementary Material: Fig. VIII). Looking in more detail at the distribution of trait levels among species, the freshwater marsh contained slightly more macropterous species, indicating a greater potential for dispersal. Species in the freshwater marsh also preferred higher moisture levels, whereas species in the saltwater marsh even included some with (steno-)xerophilic preferences. Additionally, a greater proportion of species in the saltwater marsh were endogeic (i.e., occurring below ground), indicating increased adaptation for digging.

**Figure 8:**
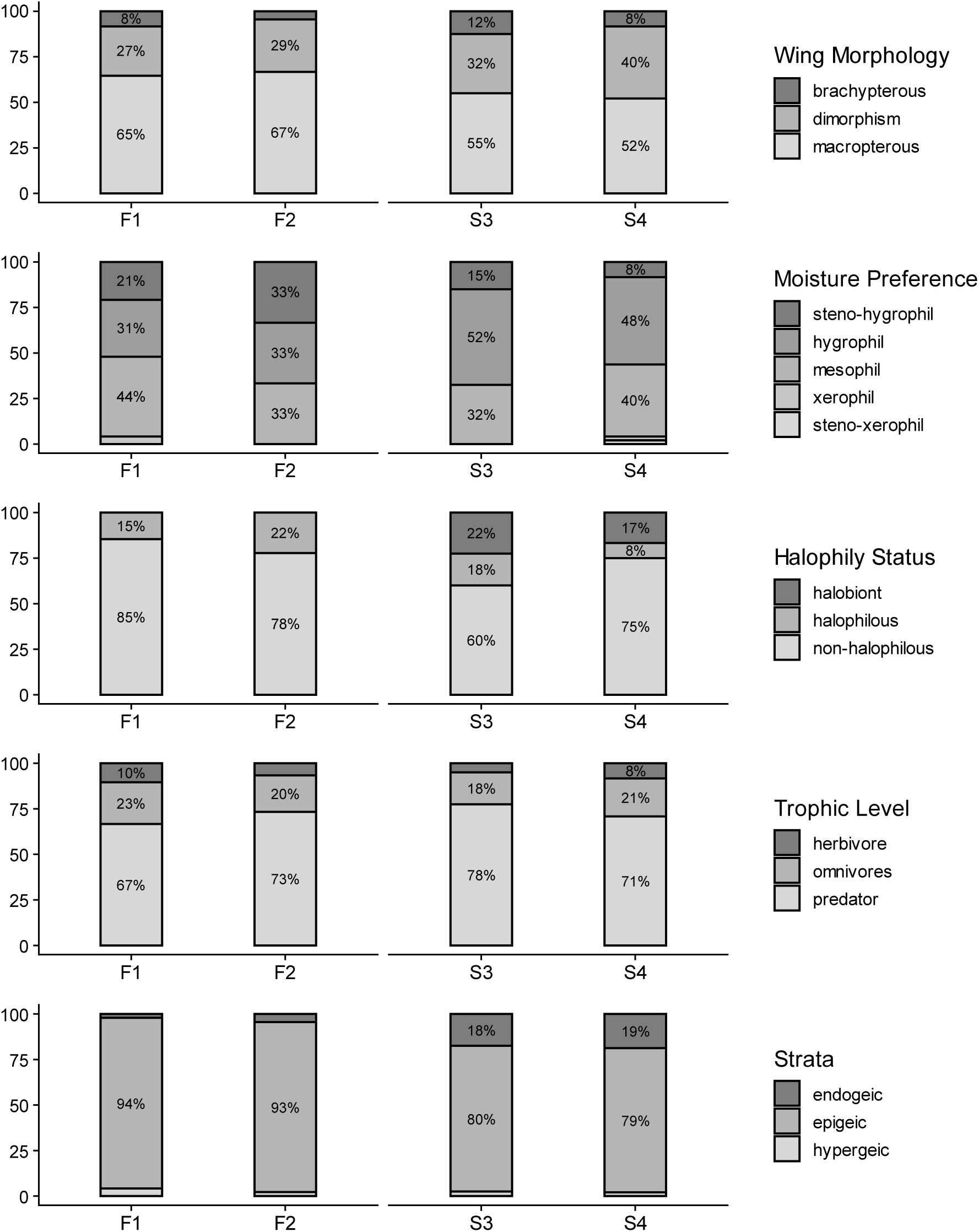
Percentage distribution of species traits (see legend) for each of the four transects. Transects F1 (N = 47) and F2 (N = 46) are located in the freshwater marsh, while transects S3 (N = 41) and S4 (N = 56) are located in the saltwater marsh of the Elbe estuary. N indicates the total number of species recorded per transect.

### Model

The final model indicated that temporal variation and functional diversity, particularly those related to habitat breadth and wing morphology, are primary drivers of carabid beetle abundance in the Elbe marshlands. Random effects accounted for variability among species and traps, with interspecies variation explaining considerable variation in activity abundance.

Specifically, the model revealed significant effects of month (χ² = 132.72, df = 6, p < 0.001), functional richness for both dispersal traits (FRic.disp: χ² = 10.32, df = 1, p = 0.001) and preference traits (FRic.pref: χ² = 6.08, df = 1, p = 0.014), functional evenness for dispersal traits (FEve.disp: χ² = 7.77, df = 1, p = 0.005), and body size (χ² = 13.87, df = 1, p < 0.001) on mean activity abundance. Marsh type did not have a significant effect (χ² = 1.10, df = 1, p = 0.295), but was retained for design consistency and due to its ecological relevance. See Table 3 for full model results. Functional richness of dispersal traits (FRic.disp: β = 11.99, p = 0.001) and preference traits (Fric.pref: β = 0.08, p = 0.014) were both positively associated with mean activity abundance, while functional evenness of dispersal traits showed a negative relationship (β = −2.65, p = 0.005). Mean body size also exhibited a positive effect (β = 0.18, p < 0.001).

**Table 3:** Results of the Generalized Linear Mixed Model predicting activity abundance of carabid beetles in the Elbe marshlands. Presented are estimates, standard errors, z-values and associated p-values for each fixed effect and variance and standard deviation for random effects, with the number of levels for each grouping factor.

| *Fixed effects* | *Estimate* | *Std. Error* | *z - value* | *Pr (>\| z\| )* |  |
| --- | --- | --- | --- | --- | --- |
| *(Intercept)* | -5.108 | 0.889 | -5.742 | < 0.001 | *** |
| *marsh_saltwater marsh* | 0.145 | 0.138 | 1.047 | 0.2949 |  |
| *month_04* | 0.399 | 0.108 | 3.679 | < 0.001 | *** |
| *month_05* | 1.002 | 0.106 | 9.455 | < 0.001 | *** |
| *month_06* | 0.649 | 0.108 | 5.981 | < 0.001 | *** |
| *month_07* | 0.481 | 0.110 | 4.33 | < 0.001 | *** |
| *month_08* | 0.021 | 0.117 | 0.175 | 0.8606 |  |
| *month_09* | 0.426 | 0.114 | 3.717 | < 0.001 | *** |
| *FRic.disp* | 11.985 | 3.730 | 3.213 | 0.0013 | ** |
| *FEve.disp* | -2.651 | 0.950 | -2.787 | 0.0053 | ** |
| *FRic.pref* | 0.075 | 0.030 | 2.466 | 0.0136 | * |
| *size_mean* | 0.176 | 0.047 | 3.724 | < 0.001 | *** |

| *Random effects* | *Variance* | *Std. Dev.* | *groups* |
| --- | --- | --- | --- |
| *species* | 3.635 | 1.907 | 84 |
| *trap* | 0.082 | 0.286 | 44 |

## Discussion

### Estuarine marshes as ecotones

Coastal marshes and estuaries are ecologically critical transition zones where terrestrial and marine influences intersect. Owing to their high productivity, capacity for carbon sequestration, nutrient cycling, and flood mitigation, these habitats provide essential ecosystem services while supporting rich biodiversity (Butzek, 2014; Greenberg et al., 2006). Species richness, community composition and functional diversity vary strongly in these marshes along environmental gradients, primarily driven by salinity and flooding regimes (Rankin et al., 2023). Our study of the Elbe estuary confirmed these global patterns, showing that freshwater and saltwater marshes contain distinct carabid beetle assemblages shaped by differing environmental stressors (Lambeets et al., 2008). The freshwater marshes harboured a higher species and functional trait richness, yet with greater dominance by generalists. In contrast, saltwater marshes, exposed to harsher osmotic and hydrologic conditions supported fewer species but higher functional evenness and divergence, indicative of niche specialization and adaptation to saline stress (De Battisti, 2021).

### Salinity, flooding regimes and trait filtering

Salinity and flooding regimes strongly influence carabid beetle assemblages by filtering species based on physiological tolerance and ecological traits (Pétillon et al., 2008). Consistent with the zonation patterns documented in the Elbe estuary, there is a pronounced community turnover along the salinity gradient (Neiske et al., 2025). Trait diversity, including attributes such as dispersal ability, moisture preference and trophic position, critically determines ecosystem resilience, particularly under cyclical disturbance and rapid environmental change, beyond taxonomic richness alone. This aligns with recent findings that functional trait distributions tend to narrow and specialize as environmental stress intensifies (Fournier et al., 2015; Litavsky et al., 2021).

Carabid assemblages in both marsh types predominantly consisted of wetland-associated species, affirming their adaptation to hydric conditions. Yet, overlap with forest and open biotope specialists indicates a notable degree of habitat generalism. Trait-level differences between marsh types further underscore the influence of hydrological and salinity gradients. Freshwater marsh species typically feature larger body sizes and higher proportions of macropterous (winged) forms with generally high moisture preferences, suggesting differences in dispersal and life-history strategies shaped by local environmental conditions (Deppe & Fischer, 2023). Salt marsh communities are dominated by halophilic and hygrophilic species, including some endogeic carabids, all specialized to cope with periodic inundation and high salinity soils. Moreover, some xerophilic species also occur, likely adapted to microhabitats representing heterogeneous conditions with limited surface moisture between inundation events (Coccia & Fariña, 2019; Yin et al., 2018). These xerophilic species may endure higher temperature and osmotic stresses, filling ecological niches distinct from the strictly halophilic and hygrophilic species assemblages (Alford et al., 2023). These findings are consistent with previous reports of marshland specialist carabids being filtered by both flooding frequency and salinity (Ridel et al., 2021; Lambeets et al., 2008; Irmler et al., 2002).

### Functional diversity’s role in resilience and ecosystem functioning

Divergent patters in functional diversity indices illustrate differing community assembly strategies across marsh types. Saltwater marshes exhibit high levels of functional richness, evenness and dispersion for preference traits, indicating pronounced trait complementarity, ecological strategies and niche differentiation, as well as a broader spectrum of roles, despite a slightly reduced species pool. In this demanding habitat the pattern is consistent with the hypothesis that suggests such trait complementarity buffers ecosystem functions against extreme salinity fluctuations, tidal inundation, and episodic disturbances, enhancing community resilience typical of coastal wetlands (De Battisti, 2021; Yeager & Hughes, 2025). Conversely, freshwater marshes exhibited higher overall functional richness but less pronounced divergence, indicative of broader, less specialized trait coverage. Carabid assemblages in these marshes are characterized by higher moisture preferences and increased dispersal potential via macroptery, reflecting lower osmotic stress and broader colonization opportunities (Den Boer, 1971). This indicates reduced abiotic filtering, permitting coexistence of a wider range of generalist and moderately specialized species (Arruda Almeida et al., 2018). While the greater functional richness in freshwater marshes may provide ecological plasticity and adaptability to a wider range of conditions, it may also reduce resilience to rapid directional environmental change or novel stressors (Song et al., 2024).

Our results demonstrate that functional richness in both dispersal- and preference-based traits positively affects carabid activity abundance, directly linking trait diversity to ecosystem processes such as nutrient cycling and food web structures. Communities with wider trait ranges can exploit broader resources, maintain more effective pest regulation, and ensure greater stability to marshland food webs.

### Beta Diversity, Turnover, Spatial Heterogeneity

High beta diversity both within and between marshes illustrates the crucial role of landscape-and environmental heterogeneity in driving regional biodiversity of carabid beetles (Mori et al., 2018). Despite modest differences in total species richness, the underlying community composition and functional traits differed considerably. Within-marsh turnover likely reflects fine scale niche partitioning, whereas the pronounced differentiation between freshwater and saltwater marshes indicates broader environmental filtering and the presence of habitat-specific assemblages. No overall gradient in activity abundance was detected with increasing distance from the riverbank. However, patterns of species-specific microhabitat use became evident, reflecting strong specialisation. These specialised species may perform distinct ecological functions along the hydrological gradient, contributing to the heterogeneous landscape of ecosystem services within the marshes. This pattern is consistent with findings from spatially extensive studies showing considerable community change, even over short distances when crossing hydrological or environmental chemical boundaries (Irmler et al., 2002). Such observations further support previous work showing that estuarine gradients, from freshwater to highly saline tidal zones, create strong environmental filters that drive species turnover and community assembly (Lambeets et al., 2008; Greenberg et al., 2006).

### Marshes as refugia and implications for conservation

Marshes serve as vital refugia for specialist and rare species, even when fragmented or isolated, supporting taxa of conservation concern with narrow habitat requirements, such as the saltmarsh specialists *Cillenus lateralis* and *Dyschirius impunctipennis*, which depend on periodic saline inundation (Topp, 1979). Certain species, like *Pogonus chalceus* and *Dyschirius chalceus* are highly disturbance sensitive and have very restricted distributions strongly tied to tidal riverbanks (Desender & Maelfait, 1999; Kotze et al., 2011). The predominance of macropterous species and wing dimorphism among marsh carabids reflects their capacity to recolonize restored habitats or populate new areas, while brachypterous forms tend to dominate stable environments (Den Boer, 1971). Despite this dispersal capacity, empirical evidence indicates that even restored marsh systems rarely replicate the full species richness or ecological functions of intact marsh ecosystems.

Small or isolated tidal marshes hold significant conservation value in the face of ongoing habitat modification and climate change and their layered spatial heterogeneity is essential for maintaining marsh ecosystem multifunctionality. This underscores the importance of protecting functionally intact marshes and guiding restoration efforts to conserve marshland systems across all hydrological zones, rather than focusing solely maximizing area and connectivity, thereby sustaining critical ecosystem functions and resilience (Pétillon et al., 2023; Veldkornet & Adams, 2025; Verschoor et al., 2025).

### Phenology and seasonal activity patterns

Seasonal activity patterns observed in this study, with earlier and bimodal activity abundance peaks in the freshwater marsh contrasted by later, more gradual increases in saltwater marsh - communities, reflect species-specific differences in phenology and reproductive phases. A closer examination of the three most abundant species in each marsh reveals that overall monthly activity patterns are strongly influenced by these dominant species (Supplementary Materials: Fig. III:). In the freshwater marsh, the activity peaks closely mirror those of the spring-breeding species *Limodromus assimilis*, which was by far the most abundant species (Supplementary Materials: Table I). These peaks correspond to its reproductive period in spring and the emergence of the next generation in autumn. Similarly, in the saltwater marsh, the second peak in overall activity abundance is mainly shaped by *Pterostichus niger*, an autumn breeder that occurred in high numbers. The earlier activity peaks observed in the freshwater marsh also reflect the breeding behaviour of its species assemblages. Approximately 70% of the species occurring in the freshwater marsh are spring breeders, while around 13% are summer or autumn breeders. In contrast, in the saltwater marsh, only about 50% of the species are spring breeders, whereas approximately 35% are summer or autumn breeders.

Furthermore, these patterns may also reflect variations in microclimate, flooding regimes, and resource availability between these marshes, all of which can further shape community dynamics (Kotze et al., 2011). The ability of carabid beetles to adjust their activity in response to microclimatic and hydrological cues emphasizes their importance in shaping trophic interactions and their value as biotic indicators for monitoring the conditions and disturbances across different marsh types. (Desender et al., 2007; Middendorf et al., 2025).

### Conclusion

This study provides an ecological baseline showing how salinity gradients and flooding regimes influence carabid beetle diversity and function. Maintaining both functional and taxonomic diversity, as well as microhabitat complexity is essential for marsh ecosystem resilience in the context of sea-level rise, increasing salinity intrusion, flood regime alterations, and anthropogenic impacts. Freshwater and saltwater marshes support distinct, trait-diverse carabid assemblages with implications for ecosystem services like nutrient cycling in these vulnerable ecotones. Conservation strategies should emphasize trait-based, long-term monitoring and multitrophic functional diversity approaches to reveal shifts in ecosystem functions, not detectable by species richness alone (Sperandii et al., 2025). Comparative studies across broader geographic regions and under different management systems are needed.

## Acknowledgements

We would like to express our gratitude to the environmental protection offices of the District of Pinneberg and the District of Steinburg as well as the Stiftung Naturschutz Schleswig-Holstein and the national park administration of the Landesbetrieb für Küstenschutz, Nationalpark und Meeresschutz Schleswig-Holstein for supporting our scientific work in the nature reserve „Haseldorfer Binnenelbe mit Elbvorland“, the Fauna-Flora-Habitat Area DE 2393-392 “Schleswig-Holsteinisches Elbästuar mit angrenzenden Flächen”, the bird protection area 2323-401 “Unterelbe bis Wedel”, and protection zone 1 of the “Nationalpark Schleswig-Holsteinisches Wattenmeer”, respectively. We thank Arne Wulff and Janina Bellmann for their assistance in the field and laboratory, as well as for their technical support. We are also thankful to Stephan Gürlich for dedicating his time to validate taxonomic identifications.

## Data availability

The datasets are available in the ZFDM Repository from the University Hamburg. [https://doi.org/10.25592/uhhfdm.18072]

