## Supplementary Material for "Diversity and Functional Roles of Carabid Beetles across Salinity Gradients in Marshlands"

### Supplementary Materials

*Species accumulation curves:*

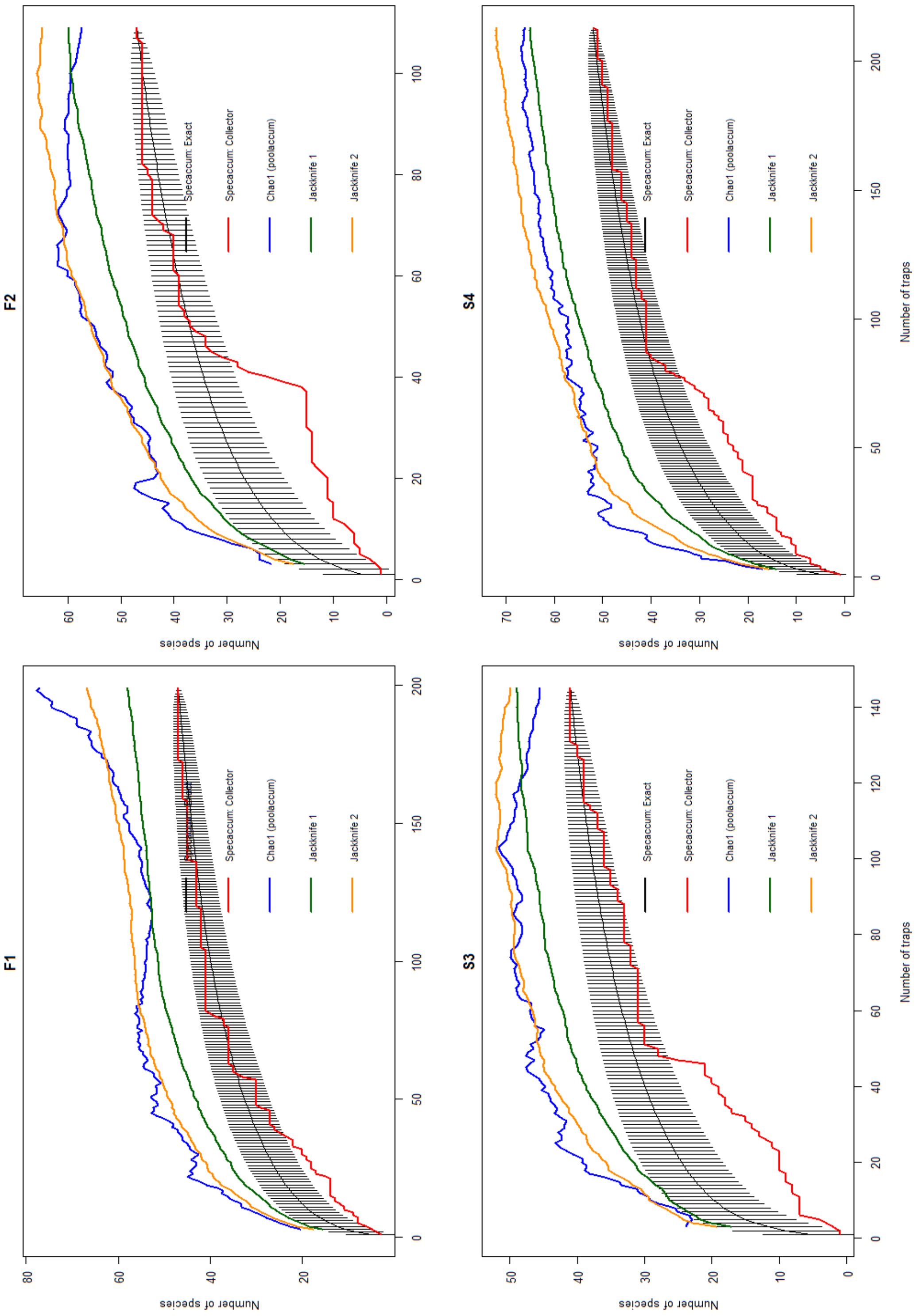


Figure I: Species accumulation (exact and collector method) and richness estimators (Chao1, Jacknife1 and Jacknife2) for carabid beetle assemblages across transects in the Elbe estuary. The x-axis shows the sampling effort (here: number of traps) and the y-axis shows the observed and estimated species richness

*Species Richness Estimator:*

The "Chao1" estimation method is described by the following equation:

$$S_P = S_O + \frac{a1^2}{2\,a2}$$

($S_P$ = estimated species richness; $S_O$ = number of observed species; a1 = number of species that occurred only once in a sample; a2 = number of species that occurred exactly twice in a sample (Gardener, 2014)).

The principle behind this equation is that, when estimating the actual number of species in a habitat, if a species is detected only once during the investigation, there is a probability that other rare species present in the habitat were not detected in the samples. As soon as species have been detected at least twice in a sample, according to the Chao1 estimation method, it is unlikely that there are additional species present in the habitat that have simply not been captured in the samples (Chao, 1987). The abundance-based coverage "ACE" estimation method is described by the following equation:

$$S_P = S_{abund} + \frac{S_{rare}}{C_{ACE}} + \frac{a1}{C_{ACE}} \times \gamma^2$$

$$C_{ACE} = 1 - \frac{a1}{N_{rare}}$$

($S_P$ = estimated species richness; $S_{abund}$ = species that occurred more than ten times in the samples; $S_{rare}$ = species that occurred ten times or less; a1 = number of species that occurred only once in a sample; $C_{ACE}$ = estimator for sample coverage; $\gamma^2$ = estimated coefficient of variation for a1; $N_{rare}$ = number of individuals of the rare species (Gardener, 2014)).

The "ACE" estimation method is more complex than the "Chao1" calculations, as it additionally includes an estimation of sample coverage, i.e., the proportion of the species richness composition based on the species in a sample. Furthermore, the equation is adjusted for the variation in the distribution through a coefficient of variation ($\gamma^2$), so that the standard deviation values given in addition to the estimates are lower than those of the "Chao1" estimation method (Chao et al., 2005).

*Habitat Types and Preferences:*

Species habitat preferences for the occurring carabid species were classified according to the Catalogue of Habitat Preferences (Gesellschaft für Angewandte Carabidologie e. V., 2009). The catalogue presents a general classification of first-level habitat types, followed by a more detailed subdivision into specific habitats. For our analysis, we used the first-level general classification as the basis for assigning habitat preferences. We grouped the first-level habitats into three broader categories. Species that were commonly found in three or more first-level habitat types according to the catalogue were classified as eurytopic.

*General classification*

(first-level habitat types)

1. Coastal Salt Biotopes & Marshlands

~~2. Mountain Biotopes~~

3. Shores, Banks & low vegetation

4.Vegetation-rich Wetlands

5. Wet & Moist Forests

6. Diverse Forest Biotopes

7. Dry & Open Biotopes

8. Open & Cultivated Landscapes

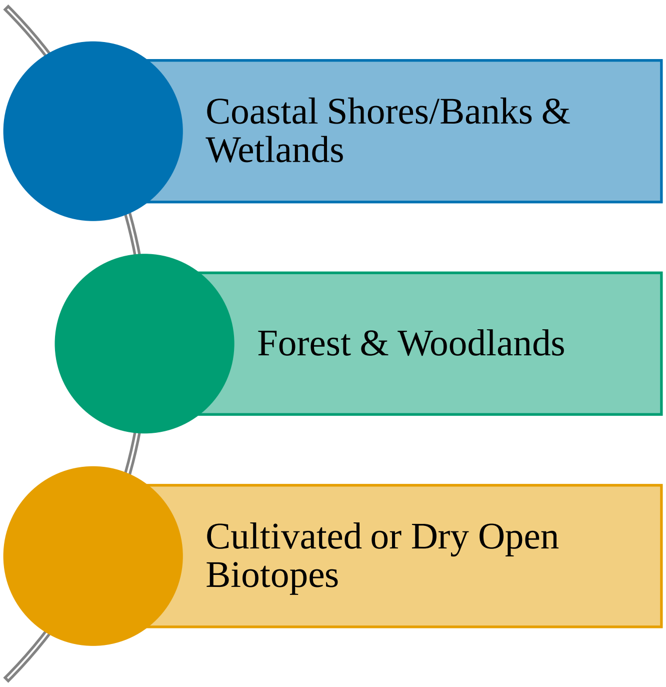

*Functional Traits:*

Prior to calculating the functional diversity indices, we assessed pairwise correlations among traits and excluded those showing high collinearity (Pearson's $|r| > 0.7$). This approach ensured a statistically robust reduction in trait dimensionality, thereby minimizing the risk of overfitting in trait space calculations.

FRic reflects the potential functional trait capacity of a community, regardless of species abundances. FEve quantifies how evenly abundances are distributed across the occupies trait space. FDiv provides insight into niche differentiation and resource use by quantifying how species abundance is distributed within the trait space of a community. It describes the extent to which the most important ecosystem functions (as reflected by abundance) are carried out by species with distinctive functional traits. Functional Dispersion (FDis) reflects how functionally dispersed or diverse the community is; high values indicate greater functional trait diversity and higher ecological niche differentiation. For this set of traits, the calculation of FDiv is not possible, since its categorial traits only. However, Functional Dispersion (FDis) can be calculated instead, using a species dissimilarity matrix.

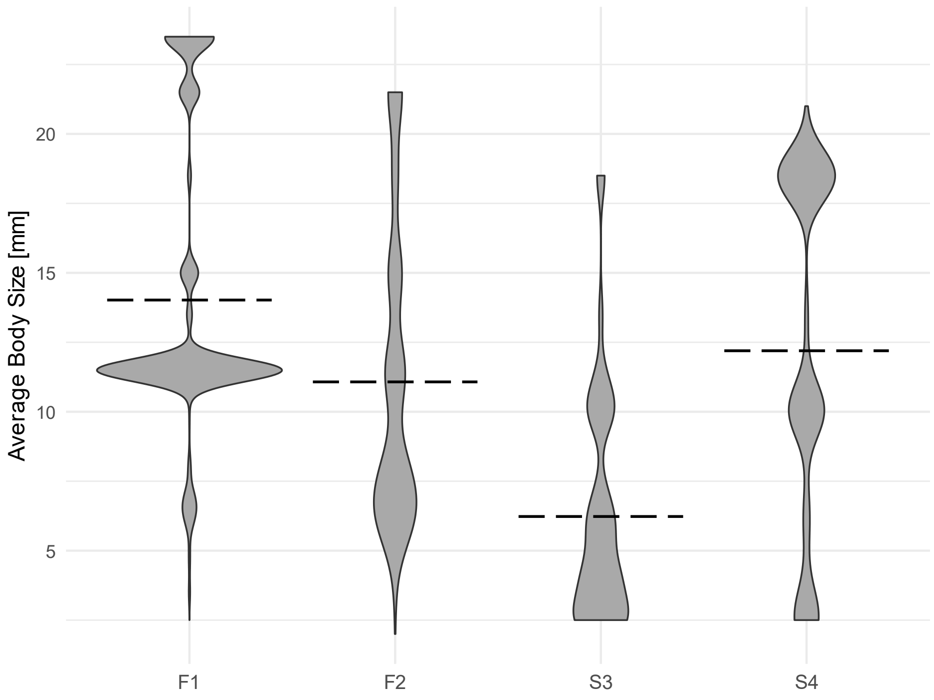


Figure I: Distribution of carabid beetle body size (mm) in freshwater (F1 & F2) and saltwater (S3 & S4) marshes visualized as violin plots. Wider sections represent a higher occurrence of individuals at those body sizes. The dashed line shows the median body size for each transect.

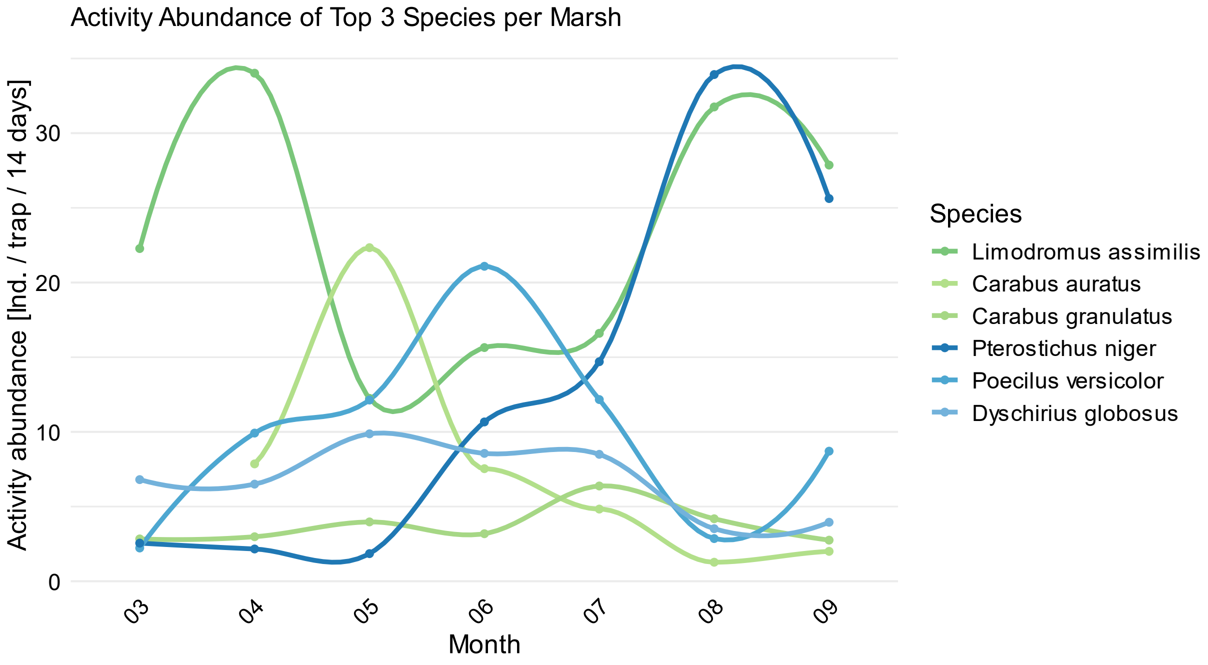


Figure II: Seasonal patterns of activity abundance of the three most abundant carabid species per marsh throughout the annual activity period type (freshwater marsh: green; saltwater marsh: blue). The lines represent LOESS curves, which use locally weighted regression to reveal non-linear trends across the months for each species.

Table I: Abundances of carabid beetle species across transects in the freshwater and saltwater marshes of the Elbe estuary. Individual counts for each species are presented for each transect (study site), summed by marsh type (freshwater or saltwater) and as overall total counts.

| Species | F1 | F2 | S3 | S4 | Freshwater | Saltwater | Total |
|---|---|---|---|---|---|---|---|
| *Acupalpus exiguus* | 0 | 1 | 0 | 0 | 1 | 0 | 1 |
| *Agonum emarginatum* | 1 | 10 | 0 | 0 | 11 | 0 | 11 |
| *Agonum marginatum* | 0 | 0 | 8 | 1 | 0 | 9 | 9 |
| *Agonum muelleri* | 0 | 1 | 0 | 0 | 1 | 0 | 1 |
| *Agonum scitulum* | 41 | 158 | 2 | 0 | 199 | 2 | 201 |
| *Agonum thoreyi* | 8 | 83 | 1 | 0 | 91 | 1 | 92 |
| *Agonum viduum* | 0 | 3 | 0 | 0 | 3 | 0 | 3 |
| *Agonum viridicupreum* | 0 | 1 | 0 | 0 | 1 | 0 | 1 |
| *Amara aenea* | 2 | 0 | 0 | 0 | 2 | 0 | 2 |
| *Amara communis* | 1 | 1 | 0 | 11 | 2 | 11 | 13 |
| *Amara convexior* | 3 | 0 | 0 | 0 | 3 | 0 | 3 |
| *Amara convexiuscula* | 0 | 0 | 2 | 9 | 0 | 11 | 11 |
| *Amara familiaris* | 4 | 2 | 6 | 1 | 6 | 7 | 13 |
| *Amara lunicollis* | 1 | 0 | 0 | 2 | 1 | 2 | 3 |
| *Amara ovata* | 1 | 0 | 0 | 0 | 1 | 0 | 1 |
| *Anchomenus dorsalis* | 183 | 1 | 0 | 0 | 184 | 0 | 184 |
| *Anisodactylus binotatus* | 1 | 7 | 0 | 1 | 8 | 1 | 9 |
| *Asaphidion flavipes* | 18 | 1 | 0 | 0 | 19 | 0 | 19 |
| *Badister bullatus* | 0 | 1 | 0 | 0 | 1 | 0 | 1 |
| *Badister lacertosus* | 1 | 0 | 0 | 0 | 1 | 0 | 1 |
| *Badister sodalis* | 4 | 0 | 1 | 92 | 4 | 93 | 97 |
| *Bembidion aeneum* | 0 | 2 | 265 | 25 | 2 | 290 | 292 |
| *Bembidion assimile* | 0 | 2 | 0 | 0 | 2 | 0 | 2 |
| *Bembidion biguttatum* | 3 | 19 | 0 | 0 | 22 | 0 | 22 |
| *Bembidion dentellum* | 1 | 2 | 10 | 2 | 3 | 12 | 15 |
| *Bembidion iricolor* | 0 | 0 | 119 | 7 | 0 | 126 | 126 |
| *Bembidion minimum* | 0 | 0 | 343 | 43 | 0 | 386 | 386 |
| *Bembidion normannum* | 0 | 0 | 16 | 4 | 0 | 20 | 20 |
| *Bembidion properans* | 4 | 1 | 96 | 117 | 5 | 213 | 218 |
| *Bembidion quadrimaculatum* | 1 | 0 | 16 | 13 | 1 | 29 | 30 |
| *Bembidion tetracolum* | 17 | 1 | 9 | 3 | 18 | 12 | 30 |
| *Bembidion varium* | 0 | 0 | 2 | 0 | 0 | 2 | 2 |
| *Blemus discus* | 0 | 1 | 0 | 12 | 1 | 12 | 13 |
| *Broscus cephalotes* | 0 | 0 | 0 | 53 | 0 | 53 | 53 |
| *Calathus erratus* | 0 | 0 | 0 | 6 | 0 | 6 | 6 |
| *Calathus fuscipes* | 0 | 0 | 1 | 5 | 0 | 6 | 6 |
| *Calathus melanocephalus* | 0 | 0 | 0 | 2 | 0 | 2 | 2 |
| *Carabus auratus* | 943 | 0 | 0 | 0 | 943 | 0 | 943 |
| *Carabus granulatus* | 418 | 159 | 0 | 0 | 577 | 0 | 577 |
| *Carabus nemoralis* | 155 | 0 | 0 | 0 | 155 | 0 | 155 |

| | | | | | | | |
|---|---|---|---|---|---|---|---|
| *Chlaenius nigricornis* | 26 | 7 | 0 | 0 | 33 | 0 | 33 |
| *Cillenus lateralis* | 0 | 0 | 2 | 4 | 0 | 6 | 6 |
| *Clivina fossor* | 1 | 60 | 157 | 110 | 61 | 267 | 328 |
| *Demetrias monostigma* | 0 | 2 | 0 | 0 | 2 | 0 | 2 |
| *Dicheirotrichus gustavii* | 0 | 0 | 5 | 1 | 0 | 6 | 6 |
| *Dyschirius chalceus* | 0 | 0 | 12 | 0 | 0 | 12 | 12 |
| *Dyschirius globosus* | 0 | 0 | 248 | 1075 | 0 | 1323 | 1323 |
| *Dyschirius impunctipennis* | 0 | 0 | 0 | 1 | 0 | 1 | 1 |
| *Dyschirius salinus* | 0 | 0 | 64 | 3 | 0 | 67 | 67 |
| *Dyschirius thoracius* | 0 | 0 | 5 | 2 | 0 | 7 | 7 |
| *Dyschirius tristis* | 0 | 0 | 2 | 0 | 0 | 2 | 2 |
| *Harpalus affinis* | 4 | 1 | 239 | 5 | 5 | 244 | 249 |
| *Harpalus latus* | 6 | 1 | 1 | 1 | 7 | 2 | 9 |
| *Harpalus rufipes* | 109 | 4 | 48 | 158 | 113 | 206 | 319 |
| *Leistus rufomarginatus* | 8 | 0 | 0 | 1 | 8 | 1 | 9 |
| *Limodromus assimilis* | 3427 | 190 | 0 | 0 | 3617 | 0 | 3617 |
| *Loricera pilicornis* | 107 | 105 | 108 | 38 | 212 | 146 | 358 |
| *Nebria brevicollis* | 413 | 5 | 15 | 12 | 418 | 27 | 445 |
| *Nebria salina* | 27 | 12 | 7 | 2 | 39 | 9 | 48 |
| *Notiophilus aquaticus* | 1 | 0 | 0 | 1 | 1 | 1 | 2 |
| *Notiophilus biguttatus* | 18 | 2 | 1 | 2 | 20 | 3 | 23 |
| *Odacantha melanura* | 7 | 2 | 0 | 0 | 9 | 0 | 9 |
| *Oodes helopioides* | 39 | 119 | 0 | 24 | 158 | 24 | 182 |
| *Ophonus laticollis* | 1 | 0 | 0 | 0 | 1 | 0 | 1 |
| *Ophonus rufibarbis* | 1 | 0 | 0 | 0 | 1 | 0 | 1 |
| *Panagaeus crux-major* | 0 | 0 | 0 | 8 | 0 | 8 | 8 |
| *Paradromius linearis* | 0 | 0 | 2 | 1 | 0 | 3 | 3 |
| *Paradromius longiceps* | 1 | 0 | 0 | 0 | 1 | 0 | 1 |
| *Paranchus albipes* | 4 | 14 | 0 | 0 | 18 | 0 | 18 |
| *Patrobus atrorufus* | 5 | 24 | 0 | 0 | 29 | 0 | 29 |
| *Poecilus cupreus* | 6 | 7 | 117 | 170 | 13 | 287 | 300 |
| *Poecilus versicolor* | 4 | 10 | 63 | 1756 | 14 | 1819 | 1833 |
| *Pogonus chalceus* | 0 | 0 | 93 | 3 | 0 | 96 | 96 |
| *Pterostichus macer* | 0 | 0 | 0 | 6 | 0 | 6 | 6 |
| *Pterostichus melanarius* | 369 | 155 | 1 | 1 | 524 | 2 | 526 |
| *Pterostichus minor* | 0 | 1 | 0 | 0 | 1 | 0 | 1 |
| *Pterostichus niger* | 68 | 69 | 105 | 3026 | 137 | 3131 | 3268 |
| *Pterostichus nigrita* | 0 | 14 | 0 | 0 | 14 | 0 | 14 |
| *Pterostichus oblongopunctatus* | 3 | 0 | 0 | 0 | 3 | 0 | 3 |
| *Pterostichus strenuus* | 53 | 21 | 4 | 111 | 74 | 115 | 189 |
| *Pterostichus vernalis* | 3 | 10 | 10 | 70 | 13 | 80 | 93 |
| *Stenolophus mixtus* | 0 | 6 | 0 | 0 | 6 | 0 | 6 |
| *Trechoblemus micros* | 0 | 0 | 1 | 0 | 0 | 1 | 1 |
| *Trechus obtusus* | 0 | 0 | 0 | 13 | 0 | 13 | 13 |
